# Air-liquid interface exposure of A549 human lung cells to characterize the hazard potential of a gaseous bio-hybrid fuel blend

**DOI:** 10.1101/2024.03.06.583741

**Authors:** Jonas Daniel, Ariel A. Schönberger Alvarez, Pia te Heesen, Bastian Lehrheuer, Stefan Pischinger, Henner Hollert, Martina Roß-Nickoll, Miaomiao Du

## Abstract

Gaseous and semi-volatile organic compounds emitted by the transport sector contribute to air pollution and have adverse effects on human health. To reduce harmful effects to the environment as well as to humans, renewable and sustainable bio-hybrid fuels are explored and investigated in the cluster of excellence “The Fuel Science Center” at RWTH Aachen University. However, data on the effects of bio-hybrid fuels on human health is scarce, leaving a data gap regarding their hazard potential. To help close this data gap, this study investigates potential toxic effects of a Ketone-Ester-Alcohol-Alkane (KEAA) fuel blend on A549 human lung cells. Experiments were performed using a commercially available air-liquid interface exposure system which was optimized beforehand. Then, cells were exposed at the air-liquid interface to 50-2000 ppm C_3.7_ of gaseous KEAA for 1 h. After a 24 h recovery period in the incubator metabolic activity and cytotoxicity of cells were assessed. Our data support the international occupational exposure limits of the single KEAA constituents and moreover indicate no adverse effect to A549 cells when exposed to a fuel mixture. This finding applies only to the exposure scenario tested in this study and is difficult to extrapolate to the complex *in vivo* situation.

## 1 Introduction

Gaseous and semi volatile organic compounds emitted by the transport sector contribute to air pollution and are proven to have adverse effects on human health [1–3]. For example, antioxidant response in lung cells significantly decreased after exposure to volatile emissions from a gasoline engine [4]. Both a significant decrease in cell viability and a significant increase in pro-inflammatory response were observed in lung cells exposed to volatile gasoline constituents [5, 6]. The WHO estimated that 4.2 million premature deaths worldwide were caused by ambient outdoor air pollution in 2019 [7], mainly due to cardiovascular and respiratory diseases, and cancers. Addressing this problem, one target of the UN’s Sustainable Development Goals is to substantially reduce the number of deaths and illnesses from hazardous chemicals and air contamination (target no. 3.9) [8]. To reduce harmful effects to the environment as well as to humans, renewable and sustainable bio-based alternatives to fossil fuels are explored and investigated in the cluster of excellence “The Fuel Science Center” (FSC) at RWTH Aachen University. Alternatives include *bio-hybrid* fuels made from biomass combined with CO_2_ using renewable energy, which can help reduce emissions [9]. Several studies reported adverse effects of gaseous biofuels on human lung cells [10, 11], however, data on effects of bio-hybrid fuels on human health is scarce, leaving a data gap regarding their hazard potential. According to the principle of Green Toxicology [12], a characterization of this hazard potential should be conducted in parallel to the development of and prior to manufacturing and distribution of bio-hybrid fuels itself.

To help close this data gap, this study investigates potential toxic effects of a bio-hybrid fuel on human lung cells, using the well-characterized human lung cell line A549 [13, 14] as an *in vitro* lung model. To increase both sensitivity of the lung model as well as relevance of the toxicological data, a state-of-the-art air-liquid interface (ALI) continuous flow exposure system is used [15, 16]. In the exposure system a direct interaction between cells at ALI conditions and gaseous chemicals takes place, representing the physiology and exposure of human lung cells more accurately compared to cells submerged in a culture medium [17–19]. Therefore, ALI exposure has become more popular in the last few years [20] since it is also a promising alternative to *in vivo* animal experiments, promoting the 3R principle (replacement, refinement, reduction) [15]. Achieving reliability, however, ALI exposure requires a complex technical setup and monitoring of important parameters [21–23], e.g. temperature and relative humidity, which is also highlighted in the present study.

The aim of this study was to establish an *in vitro* ALI exposure system for investigating the potential toxicity of a bio-hybrid fuel blend candidate on A549 human lung cells. A promising Ketone-Ester-Alcohol-Alkane (KEAA) gasoline-like fuel blend was chosen based on previous work [24]. The approach presented here includes the adjustment of a commercially available ALI exposure system, its application for ALI experiments as well as the generation and chemical analysis of the test gas.

## 2 Materials and methods

### 2.1 Fuel blend

The KEAA fuel blend [24] used in the present study consists of 41 mol% 3-methylbutanone, 25 mol% ethanol, 16 mol% methyl acetate, 12 mol% ethyl acetate, 4 mol% pentane and 2 mol%. All constituents (analytical grade) were purchased from VWR International GmbH and were blended in the lab.

### 2.2 ALI exposure system

The exposure system used for ALI experiments was bought from Vitrocell Systems GmbH (Waldkirch, Germany). A schematic setup of the exposure system is shown in Fig 1. The whole system includes several devices for controlling mass flow, dilution, humidity, and temperature of the test gas. First, feed gas enters the main flow pipe (stainless steel) of the exposure system by applying negative pressure using a vacuum pump (SECO SV 1008 C, Busch Vacuum Solutions, Maulburg, Germany). Then, the feed gas passes two dilutors in series (total dilution ratio up to 1:100), in which parts of the feed gas are extracted and humidified clean air is added using mass flow controllers (MFC Serie 358, ANALYT-MTC Messtechnik GmbH, Müllheim, Germany). After each dilutor, temperature and humidity of the test gas are measured. Then the test gas is transported into a VITROCELL 6/4 stainless steel exposure module (Module 1 & 2) through a bypass of chemically resistant Iso-Versinic tubes (Saint-Gobain Performance Plastics, Akron, USA). TYGON tubing (R 3603, Saint-Gobain Performance Plastics, Akron, USA) is used for most of the other connections in the system. The test gas enters the exposure modules through trumpet-shaped inlets in the lid. Inside the exposure modules, cells grown on membranes are supplied with basolateral culture medium while being exposed apically to the test gas. Cells exposed to humidified clean air in a VITROCELL 6/3 stainless steel exposure module (Clean Air) are used as a control to the test gas. More technical details on the modules can be found in a previous study [25]. The flow rate (0-20 mL/min) of the test gas across the cell monolayer is adjusted by using vacuum calibration valves connected to a second vacuum pump (Laboport N840FT.18, KNF Neuberger GmbH, Freiburg, Germany). Both exposure modules and the control module are connected to a heated water circuit (CC-106A, Peter Huber Kältemaschinenbau AG, Offenburg, Germany) to maintain a constant temperature inside the modules. Most of the devices, tubing and modules are housed in a climatic chamber. Temperature inside the climatic chamber is monitored (Pt100 type probes, testo 176 T2, testo SE & Co. KGaA, Lenzkirch, Germany), and heat is distributed by heaters and fans.

**Fig 1.**
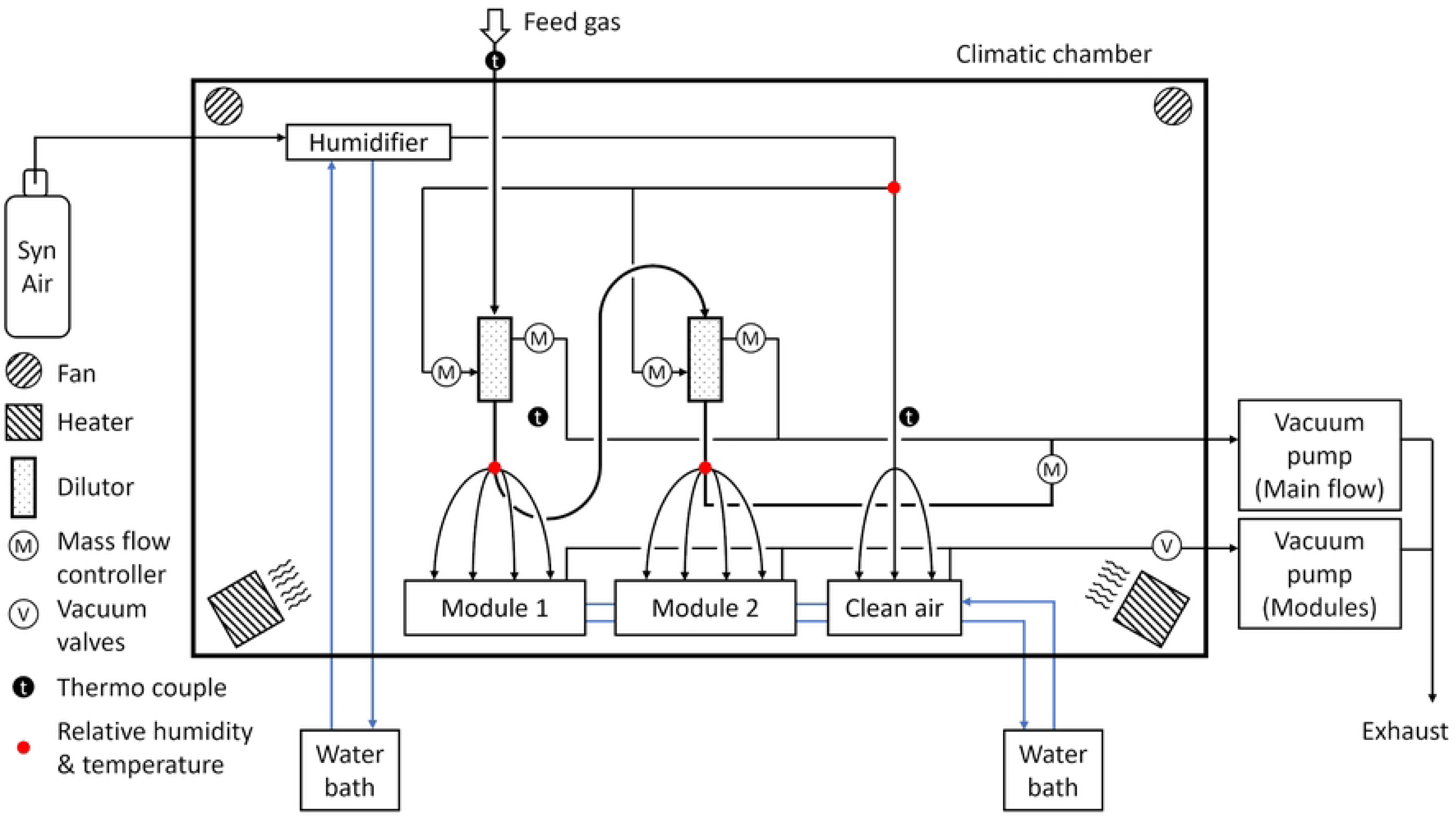
Schematic of the exposure system used to expose cells at air-liquid interface to gaseous chemicals. *(adapted from Vitrocell Systems GmbH).*

### 2.3 Adjustment of the ALI exposure system

Pre-experiments using humidified clean air as test gas were performed to validate the exposure system as well as to optimize the following experiment parameters to achieve high cell viability. In the first experiments, the heaters in the climatic chamber and the water circuit system were set to 37 °C. The position of the heaters and fans in the climatic chamber and the flow direction of the water circuit in the hull of the exposure and control modules were in ‘standard settings’, which resulted in an uneven heat distribution inside the climatic chamber as well as a temperature gradient along the exposure and control modules, increasing the risk of losing humidity from the test gas due to condensation. In this setup, the relative humidity (rH) of the test gas entering the exposure modules was *calculated* to be 68-82 % at 37 °C after humidifying the dry feed gas (N_2_) in both dilutors (1:5 ratio) with humidified clean air which was *measured* to be at 85 % rH. These conditions resulted in low cell viability in the experiments. Therefore, temperature of the climatic chamber (40 °C) and water circuit (38.5 °C), as well as humidity of the feed gas (40 g/m³) were increased to achieve high cell viability, creating incubator-like ambient conditions during exposure. To verify this, a RH/T Controller (Vitrocell Systems GmbH) was added to the exposure system and inline RH/T sensors downstream of each dilutor and upstream of the exposure modules (red dots in Fig 1) were installed to monitor (Vitrocell Monitor, Version 1.05) temperature and rH of the test gas in proximity to the cells. Moreover, heaters and fans in the climatic chamber were repositioned for better heat distribution and the flow of the water circuit for exposure and control modules was improved according to Leibrock et al. [26]. In general, these adjustments reduced loss of humidity from the test gas by lowering the risk of condensation at thermal bridges inside the exposure system, thus increasing cell viability.

### 2.4 Generation and analysis of the feed gas

The feed gas for all ALI experiments was generated on a custom-built model gas test bench (MGTB) made from grade 2 titanium (Fig 2). The feed gas is convectively heated with a closed loop control to achieve the desired temperature. All gas pipes are as well made from titanium and heated to 37 °C to prevent condensation of components from the feed gas. Various mass flow controllers (MFC) type SLA5850 (Brooks Instrument LLC, Hatfield, PA, USA) were used to regulate the flows of N_2_ (99.999 vol. %, Nippon Gas), O_2_ (99.999 vol. %, Westfalen Gas) and NO_2_ (5000 ppm in N_2_, Westfalen Gas). A HovaPOR LF-1200 (IAS GmbH, Oberursel, Germany) was used to supply water, vaporized in another part of the N_2_ flow, to humidify the feed gas. The liquid KEAA hydrocarbons were dosed with a syringe pump (Cetoni GmbH, Korbußen, Germany) and evaporated in a self-designed titanium evaporator which also uses a part of the N_2_ flow. A variety of different measuring devices was used for the gas analysis bypassing the Vitrocell exposure system: A MultiGas 2030 HS Fourier-transform infrared (FT-IR) spectroscopy gas analyzer (MKS Instruments Inc., Andover, MA, USA) for measuring hydrocarbon species (HC) and a flame-ionization detector (FID) (Thermo-FID MP by SK-Elektronik GmbH, Leverkusen, Germany) for total HC amount normalized to C_3_ and for NO_2_ was used; in serial arrangement an H_2_O condenser for measuring water content and a measurement system (FEV Europe GmbH, Aachen, Germany) containing a paramagnetic detector (PMD) for O_2_ measurement was installed. The gas matrix used was adapted to the different experiments, using the same total gas flow of 17.4 L/min (standard conditions/NIST) with a N_2_ gas flow as equilibrium.

**Fig 2.**
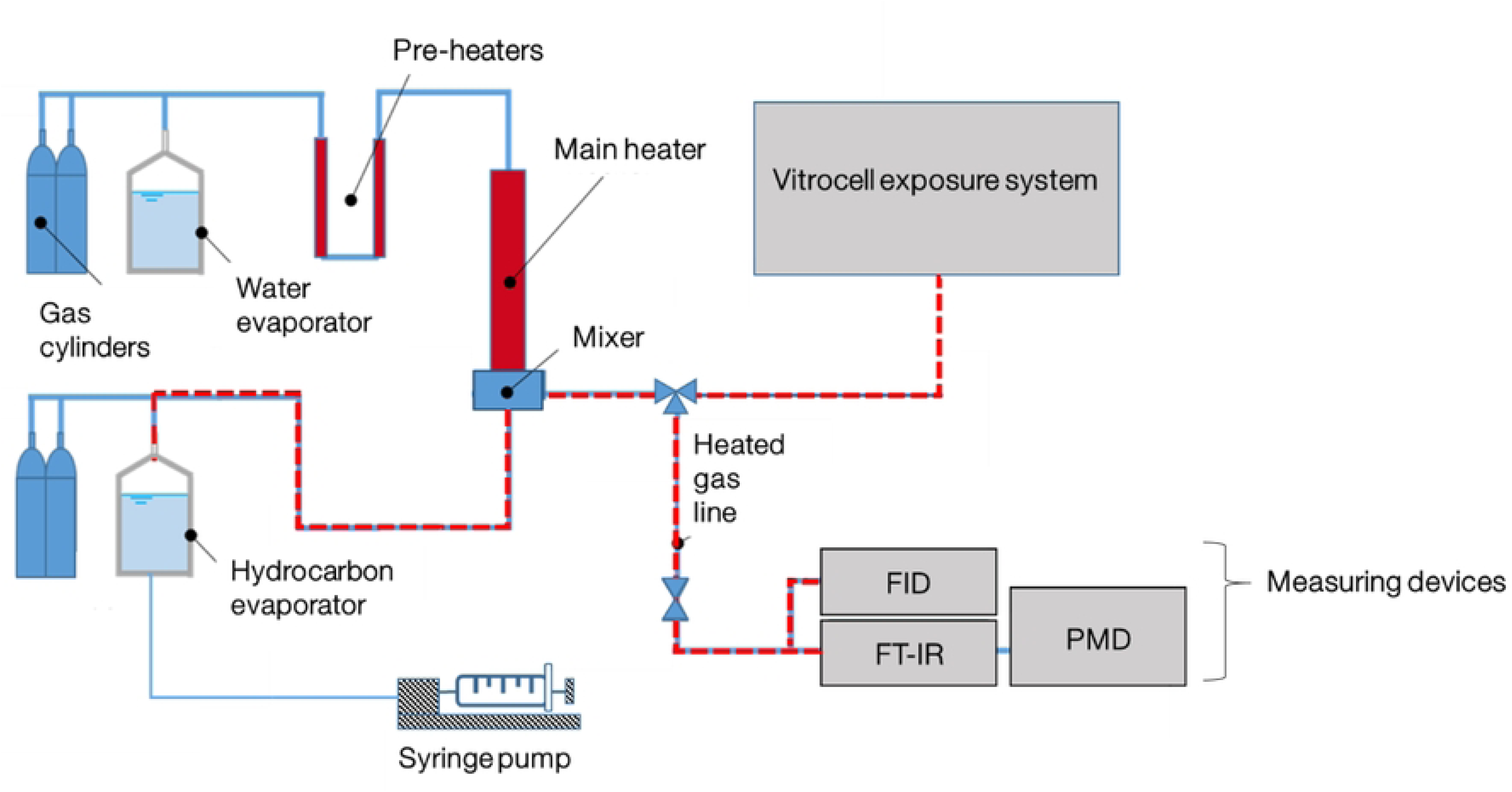
Schematic setup of the model gas test bench with the connected Vitrocell exposure system. The liquid hydrocarbons (HCs) were dosed into an evaporator using a syringe pump. Evaporated HCs were transported via a humidified gas stream (N_2_:O_2_, 80:20, % v/v) through heated pipes to the Vitrocell exposure system and measuring devices. HC species were measured using a Fourier-transform infrared (FT-IR) analyzer and for total HC amount a flame-ionization detector (FID) was used. For O_2_ measurement a paramagnetic detector (PMD) was used.

### 2.5 Dilution of the feed gas

From the total gas flow of the MGTB, a 5 L/min feed gas stream was extracted and transported to the Vitrocell exposure system. In the exposure system, the feed gas stream was diluted 1:5 with humidified clean air in both dilutors, which yields a 1:5 dilution (first dilutor) and a 1:25 dilution (second dilutor), respectively. Therefore, two concentrations of the KEAA blend were tested in parallel per experiment. In total, KEAA concentrations ranging from 50-2000 ppm C_3.7_ were tested. Concentrations of the single KEAA constituents after dilution were calculated from measured concentrations in the undiluted feed gas and are presented in Table 1.

**Table 1.** Calculated concentrations of single KEAA constituents after dilution based on measurements of the undiluted feed gas.

| Substance | Total HC concentration C<sub>3.7</sub> ppm |  |  |  |  |  |  |  |
| --- | --- | --- | --- | --- | --- | --- | --- | --- |
|  | 50 | 100 | 200 | 250 | 400 | 500 | 1000 | 2000 |
|  | ppm |  |  |  |  |  |  |  |
| 3-Methylbutanone | 15.9 | 31.8 | 63.6 | 79.5 | 127.1 | 158.9 | 317.8 | 635.6 |
| Ethanol | 9.9 | 19.8 | 39.7 | 49.6 | 79.3 | 99.2 | 198.3 | 396.6 |
| Mehtyl acetate | 6.1 | 12.3 | 24.5 | 30.7 | 49.0 | 61.3 | 122.6 | 245.2 |
| Ehtyl acetate | 4.9 | 9.8 | 19.7 | 24.6 | 39.4 | 49.2 | 98.4 | 196.8 |
| Pentane | 1.6 | 3.1 | 6.3 | 7.9 | 12.6 | 15.7 | 31.4 | 62.8 |
| Methanol | 0.9 | 1.8 | 3.5 | 4.4 | 7.1 | 8.9 | 17.7 | 35.4 |

Humidified clean air was generated from synthetic air (grade 5.0) passing a humidifier (Naphion cartridge) connected to a water bath (KISS 104A, Peter Huber Kältemaschinenbau AG, Offenburg, Germany) (Fig 1). Temperature and relative humidity of humidified clean air (37 °C, 85 % rH) were measured upstream of the dilutors using a RH/T sensor (testo 645, testo SE & Co. KGaA, Lenzkirch, Germany).

NO_2_ spiked feed gas from the MGTB was diluted 1:5 with humidified clean air in one dilutor of the exposure system to a nominal concentration of 10 ppm.

### 2.6 Cultivation of A549 cells

A549 cells were obtained from the German Collection of Microorganisms and Cell Cultures GmbH (DSMZ, ACC 107, Braunschweig, Germany) and cultured in growth medium (DMEM, Gibco 21885108, Fisher Scientific GmbH, Schwerte, Germany) supplemented with 9 % heat inactivated FBS (Gibco 10500064) at 37 °C, 95 % rH and 5 % CO_2_. Cell culture medium of subconfluent cells was aspirated, cells were rinsed with PBS without ions (Sigma-Aldrich Chemie GmbH, Taufkirchen, Germany) and detached from cell culture flasks (CytoOne, Starlab GmbH, Hamburg, Germany) using a 0.05/0.02 % Trypsin/EDTA solution (SAFC, Sigma-Aldrich). Trypsination was stopped with growth medium and cells were reseeded in fresh growth medium. For experiments, A549 cells at passages 7-20 were used.

### 2.7 Exposure procedure

#### 2.7.1 Cultivating A549 cells on inserts at ALI conditions

Forty-eight hours before exposure, A549 cells were seeded on the apical side of 6-well-sized inserts (4.524 cm² area, pore size 0.4 µm, transparent membrane, Greiner Bio-One, Frickenhausen, Germany) at a density of 60.000 cells/cm² in 1 mL growth medium (Fig 3), according to Ruth et al. [27]. Before seeding, inserts without cells were placed into microplates (CytoOne, Starlab GmbH) filled with 1.5 mL growth medium in the basal compartment and were incubated for 30 min in the incubator to condition the membranes of the inserts. After seeding, the plates containing the inserts were not moved for 5 min to let the cells sediment onto the insert membrane, supporting a homogenous cell distribution across the growth surface of the insert membrane [28]. Then, A549 cells on inserts were pre-incubated at 37 °C, 95 % rH and 5% CO_2_. After 24 h pre-incubation, basal medium in microplates was aspirated and replaced with 1.5 mL ALI medium. ALI medium consisted of DMEM without phenol red (Gibco, 11880036) with 3 % heat inactivated FBS, 25 mM HEPES (Gibco, 15630056), 4 mM GlutaMAX (Gibco, 35050038) and 100 U/mL penicillin/streptomycin (Gibco, 15140122). Medium on the apical side of the inserts was aspirated and the cell monolayer was rinsed with 1.5 mL PBS. Air-lifted inserts were pre-incubated for another 24 h until beginning of exposure.

**Fig 3.**
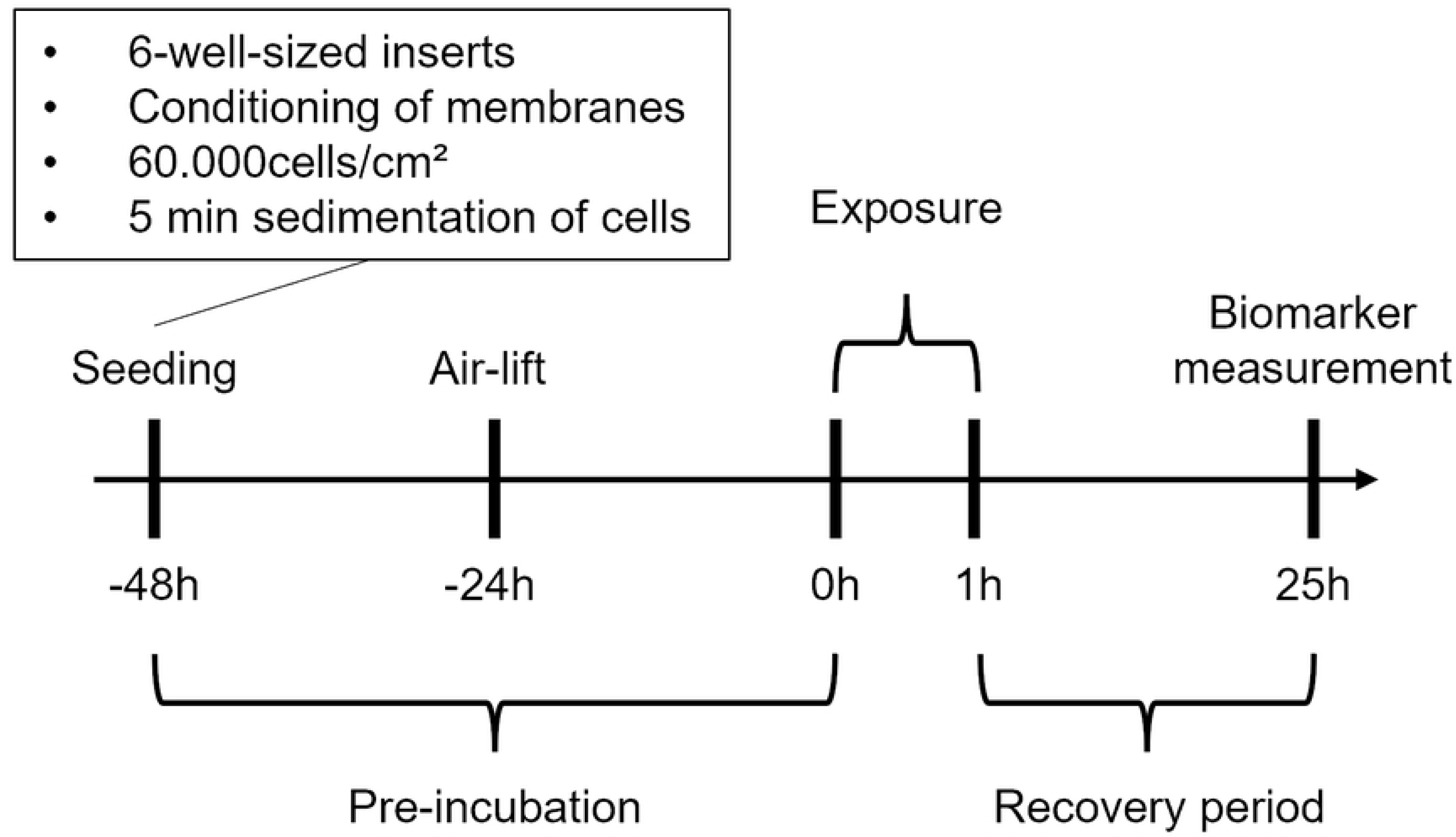
Timeline of air-liquid interface experiments.

#### 2.7.2 Calibrating the ALI exposure system for experiments

Before the start of exposure, the two clean (rinsed with 70 % ethanol) exposure modules and the control module for clean air were connected to a water bath and pre-warmed to 37° C for approx. 2 h. Then, under sterile conditions 6.7 mL and 19.7 mL of pre-warmed ALI medium were dispensed per slot in the exposure modules and in the control module, respectively. The difference in volume between exposure modules and control module was due to different cup sizes per module slot. The air-lifted inserts with A549 cells were taken out of the incubator, placed into the module slots, and checked for air bubbles below the membranes. During this step, the laminar airflow of the sterile bench was briefly turned off until the inserts with cells were enclosed within the modules, avoiding A549 cells drying out. The distance between the trumpet-shaped inlets and inserts inside the modules was 2 mm. Then, the modules were put into the climatic chamber and were connected to the water circuit. Until connection to the main flow or clean air, respectively, the inlets of the modules remained sealed with rubber plugs.

Approx. 2 h before the beginning of exposure, both the water bath for the humidifier and the water circuit for the exposure and control modules were set to 38.5 °C. The climatic chamber was heated overnight to 40 °C. The closed bypass transporting the feed gas from the MGTB to the exposure system (Fig 2) was connected to the main flow of the exposure system and was opened as the feed gas reached temperatures below 40 °C to prevent damage to the exposure system. At the same time, the main flow vacuum pump (5 L/min) and dosing of clean air were activated. Then, mass flow, pressure, temperature and rH of the main flow and clean air were calibrated using nitrogen as feed gas. The vacuum pump for the exposure and control modules was briefly activated to set the vacuum valves for the flow rate across the cells to 20 mL/min using a calibration meter (GFM 17, ANALYT-MTC Messtechnik GmbH, Müllheim, Germany). During calibration, which took approx. 1.5 h, the exposure and control modules remained disconnected from the main flow and clean air, respectively, leaving the cells undisturbed.

#### 2.7.3 Exposure of A549 cells at the ALI

As the temperature of the feed gas (N_2_) from the MGTB reached 37 °C, the inlets of the exposure and control modules were carefully connected to the main flow and clean air of the exposure system, respectively. The whole system was stabilized for approx. 15 min before exposure started. Exposure started as the vacuum pump for the exposure and control modules was activated. At the same time, dosing of gaseous KEAA, O_2_ (20 vol%) and water (40 g/m³) from the MGTB started and A549 cells were exposed at the ALI to 50-2000 ppm C_3.7_ KEAA blend for 1 h at a flow rate of 20 mL/min. Humidified clean air (37 °C, 85 % rH) was used as a negative control (clean air control).

Additionally, air-lifted cells in a microplate containing 1.5 mL basal ALI medium were incubated in the incubator to account for unspecific effects on viability of cells while being handled outside the incubator (incubator control). In separate experiments, NO_2_ (10 ppm) spiked humidified clean air was used as test gas to elicit a significant cytotoxic response from A549 cells at ALI conditions (positive control). Experiments were performed twice (n=2) in four technical replicates for each KEAA blend concentration and NO_2_ or in triplicates for clean air and incubator controls, respectively. After exposure, the dosing and gas flow were shut down by closing the bypass from the MGTB and deactivating the vacuum pumps. Then, exposure and control modules were transported back to the sterile bench. Laminar airflow was briefly turned off again to prevent damage to the cells while transferring inserts from the modules and incubator control plate to new 6-well microplates containing 1.5 mL fresh basal ALI medium. Then, cells were post-incubated under ALI conditions for 24 h in the incubator.

### 2.8 Metabolic activity and LDH release assay

After a 24 h recovery period in the incubator at ALI (Fig 3) metabolic activity and lactate dehydrogenase (LDH) release, as a marker for cell viability and necrosis of the exposed cells were assessed according to the manufacturer’s instructions. Briefly, an 1X alamarBlue HS Cell Viability Reagent (Life Technologies Corporation, Eugene, USA) solution was prepared in ALI medium. Inserts were rinsed with 1.5 mL PBS and 1 mL reagent solution was added apically to the cells. Then, inserts were transferred to new 6-well microplates without basal medium and incubated for 1 h in the incubator. After incubation, the supernatant reagent solution was homogenized and 100 µL supernatant per insert were transferred to a 96-well plate (CytoOne, Starlab GmbH) in duplicates. Fluorescence of supernatants was measured at 555/596 nm excitation/emission wavelength (Cytation 5 Multi-Mode Microplate Reader, Agilent, Waldbronn, Germany). 1X alamarBlue reagent solution in ALI medium was used as blank.

For investigating cytotoxicity of A549 cells, LDH release into the basolateral medium was measured using CyQUANT LDH Cytotoxicity Assay Kit (Life Technologies Corporation) according to manufacturer’s instructions. After the recovery period, 50 µL of homogenized basolateral medium per insert were transferred to a 96-well plate in duplicates and 50 µL of LDH assay reagent mixture were added. After 30 min incubation at room temperature in the dark, absorbance of samples was measured at 490 nm with a reference wavelength of 680 nm (Cytation 5). As an assay positive control, unexposed cells incubated at ALI conditions were lysed apically with 1.5 mL 1 % Triton X-100 (TX1) 1 h before end of recovery period. After 1 h, supernatant medium was tested for LDH release as described above. ALI medium was used as blank.

### 2.9 Statistical analysis

Raw fluorescence and absorbance data were blanked, normalized, and expressed as percentage metabolic activity of clean air control or percentage of released LDH of assay positive control (TX1), respectively. To detect significant differences between controls and treatments, ToxRat Professional (Version 3.3.0, ToxRat Solutions GmbH, Alsdorf, Germany) was used. Depending on normality (Shapiro-Wilk test) and variance homogeneity (Levene test) of data, multiple comparisons of treatments against the clean air control (multiple sequentially-rejective Welsh-t-test after Bonferroni-Holm) or two sample comparisons between controls (Welch t-test or Kolmogorov-Smirnov test) was performed. Statistical significance was indicated as *p<0.05, **p<0.01, ***p<0.001 and ****p<0.0001. Figures were created with GraphPad Prism (Version 6.07, GraphPad Software, Inc.).

## 3 Results

### 3.1 Physico-chemical analysis of feed gas and test gas

In Fig 4, a representative total hydrocarbon (HC) concentration (normalized to C_3.7_) of the dosed KEAA blend in the feed gas is shown. Concentration (5000 C_3.7_ ppm) and temperature (37 °C) of the feed gas remained constant during exposure. Before (calibration) and after exposure (shutdown), additional checks confirmed constant HC dosing.

**Fig 4.**
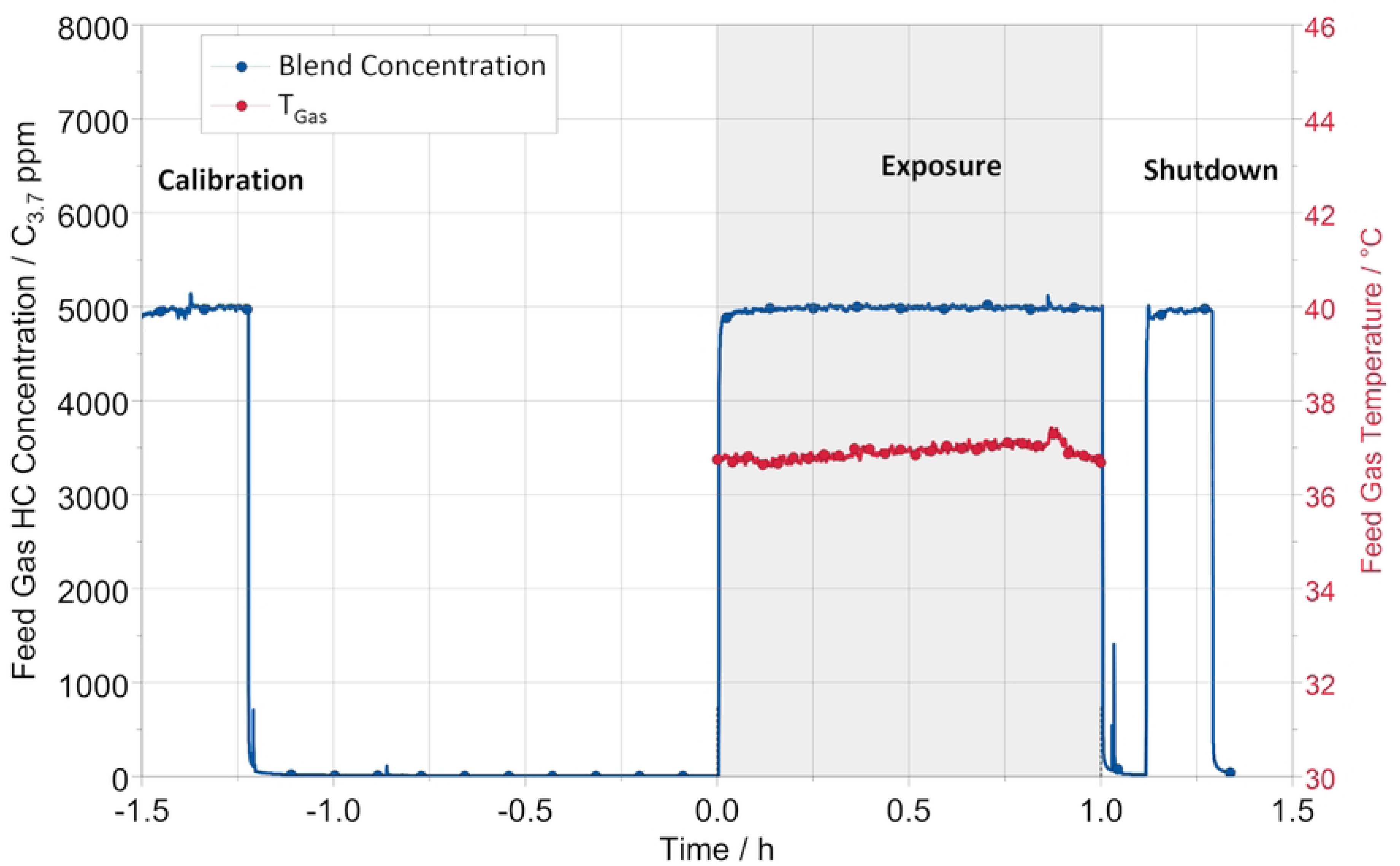
Representative feed gas concentration curve during the exposure procedure. The blue curve represents the blend concentration (KEAA) normalized to C_3.7_ in the feed gas before dilution in the exposure system. The red curve represents the feed gas temperature during the exposure phase, which is indicated by the gray background. Before (calibration) and after exposure (shutdown), the concentration level was additionally checked.

In Fig 5, representative single concentrations of KEAA blend constituents in the test gas are shown for a total HC concentration of 1000 C_3.7_ ppm (1:5 dilution of feed gas containing 5000 C_3.7_ ppm KEAA). Calculated concentrations in the exposure module were constant during exposure and as follows: 3-methylbutanone (317.8 ppm), ethanol (198.3 ppm), methyl acetate (122.6 ppm), ethylacetate (98.4 ppm), pentane (31.4 ppm) and methanol (17.7 ppm).

**Fig 5.**
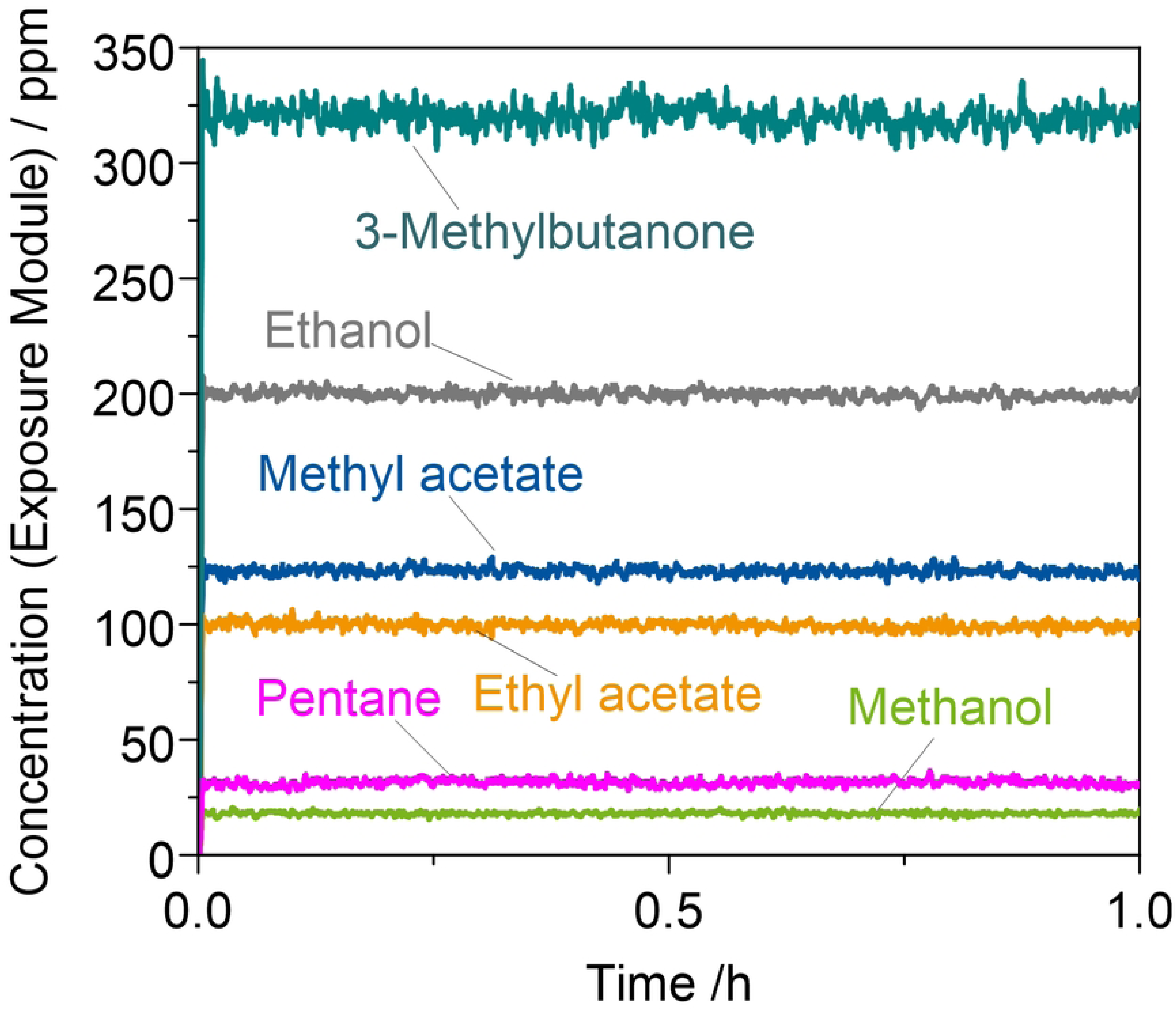
Representative calculated concentrations of single KEAA constituents after dilution of the feed gas during exposure.

Temperature and relative humidity (rH) of the test gas upstream of the two exposure modules were monitored during experiments (Fig 6). Before exposure (calibration), temperature and rH fluctuated due to opening of the climatic chamber to connect the exposure and control modules. During exposure, temperature and rH were constant at approx. 38 °C for both modules and 60 and 66 % rH for module 1 (M1) and module 2 (M2), respectively. After exposure (shutdown), temperature and rH dropped.

**Fig 6.**
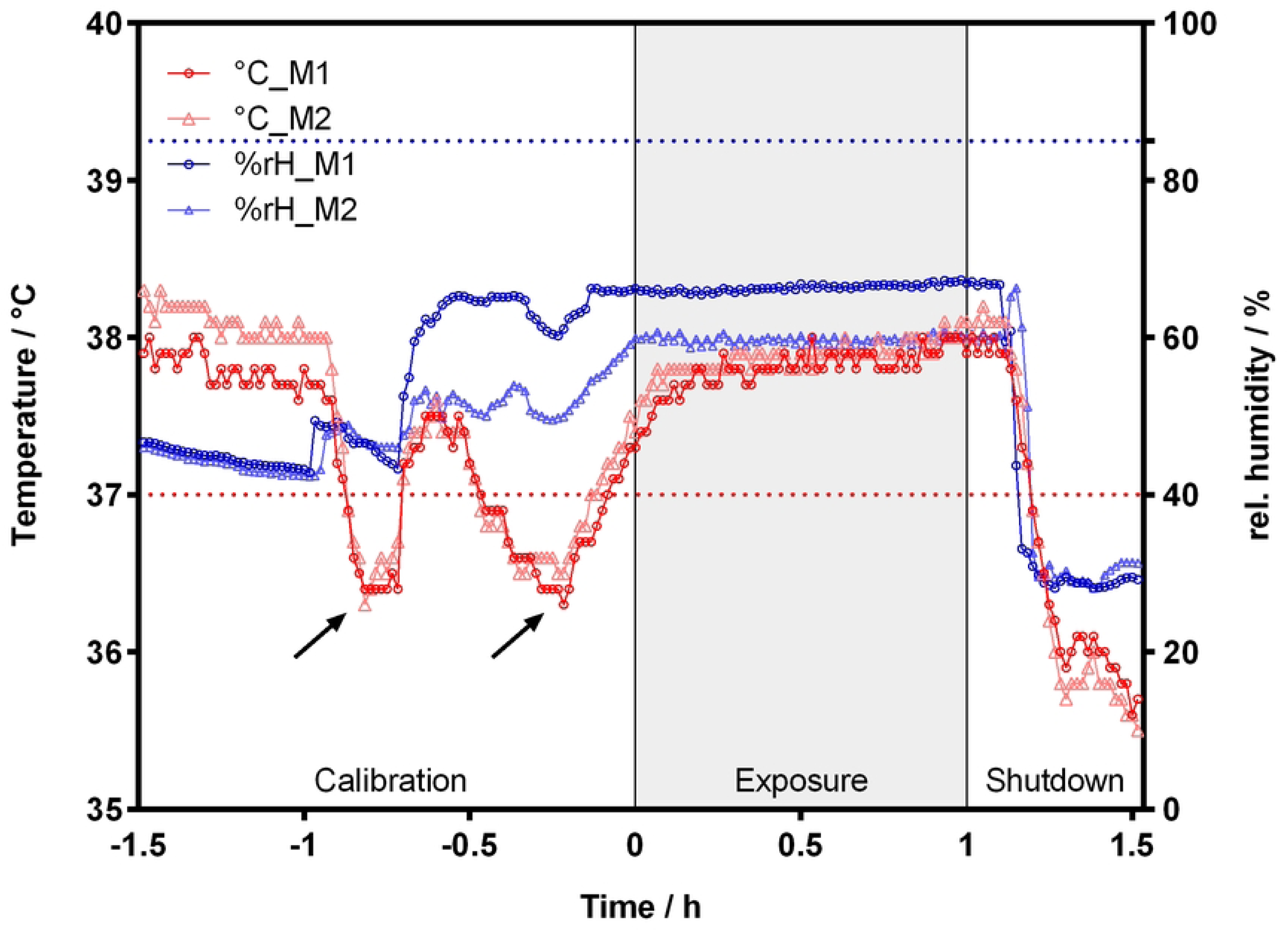
Temperature and relative humidity of the test gas during experiments. The temperature (red) and relative humidity (blue) of the test gas upstream of exposure module 1 (M1) and exposure module 2 (M2) were measured. The dotted horizontal lines indicate 37 °C (red) and 85 % rH (blue). Arrows mark the opening of the climatic chamber to insert exposure and control modules. Each experiment was divided into three phases: calibration, exposure, and shutdown.

### 3.2 Metabolic activity

As shown in Fig 7, metabolic activity in A549 cells exposed to 10 ppm NO_2_ was significantly lower compared to the clean air control (CA) (17.8 ± 5.3 %). The incubator control (IC) showed significantly higher metabolic activity (111.7 ± 3.4 %) than the CA. Metabolic activity of cells treated with KEAA was significantly different from the CA for the 500 ppm C_3.7_ treatment only (94.6 ± 5.6 %). However, no trend for concentration-dependent change in metabolic activity was observed.

**Fig 7.**
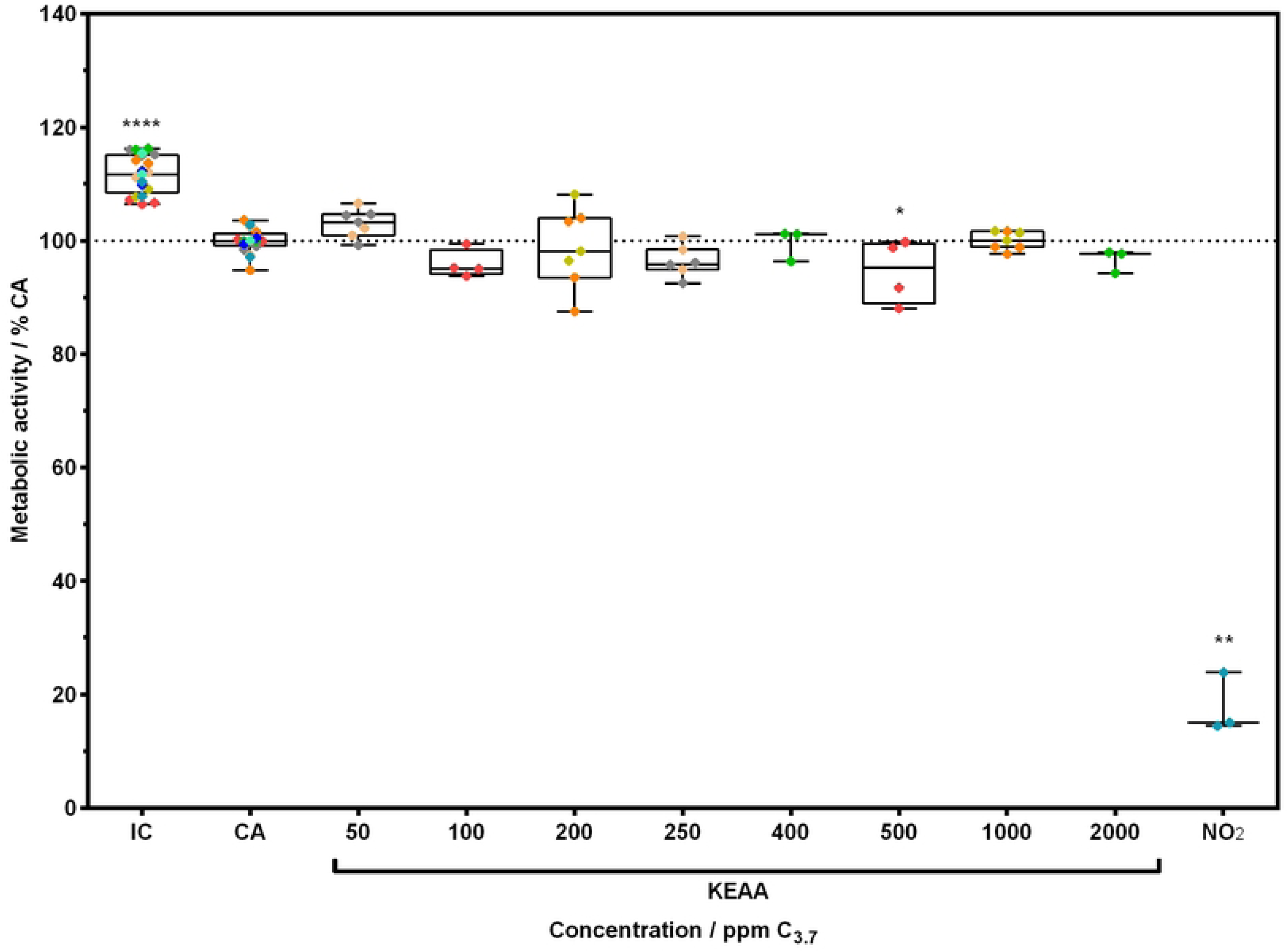
Metabolic activity of A549 cells after exposure to gaseous KEAA fuel blend. Cells at ALI conditions were exposed to humidified clean air spiked with different concentrations of KEAA or NO_2_ (10 ppm) for 1 h at a flow rate of 20 mL/min. After a 24 h recovery period at ALI conditions in the incubator, metabolic activity was investigated. Data was normalized and compared to clean air control (CA). Incubator control (IC) shows unexposed cells. Colored diamonds show technical replicates from independent experiments where one color refers to a set of two concentrations and controls.

### 3.3 LDH release

Cells exposed to 50, 250, 500 and 1000 ppm C_3.7_ KEAA showed significantly higher cytotoxicity as measured as LDH release (5.1 ± 0.8 %; 7.9 ± 1.3 %, 5.8 ± 2.6 % and 5.3 ± 2 %, respectively) compared to the CA (Fig 8). LDH release of the CA was 2.6 ± 1.4 % of the assay positive control (TX1). Cells treated with TX1 and NO_2_ showed a significant increase in LDH release compared to the CA (100 ± 16.6 % and 26.1 ± 7 %, respectively). The IC showed significantly lower LDH release (1.9 ± 2.6 %) than the CA.

**Fig 8.**
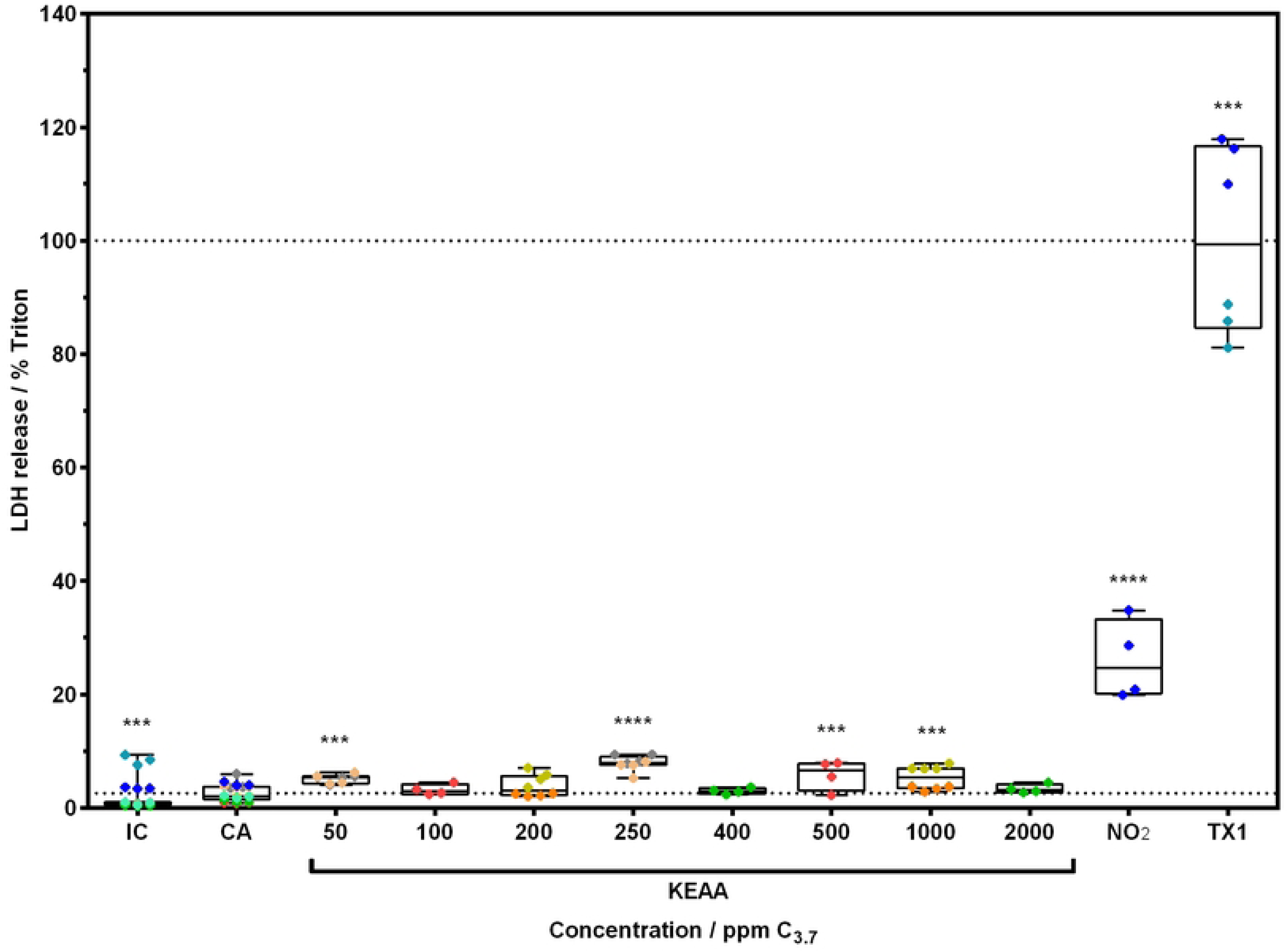
LDH release of A549 cells after exposure to gaseous KEAA fuel blend. Cells at ALI conditions were exposed to humidified clean air spiked with different concentrations of KEAA or NO_2_ (10 ppm) for 1 h at a flow rate of 20 mL/min. After a 24 h recovery period at ALI conditions in the incubator, LDH release was investigated. Data was normalized to assay positive control (TX1) and compared to clean air control (CA). Incubator control (IC) shows unexposed cells. Colored diamonds show technical replicates from independent experiments where one color refers to a set of two concentrations and controls. Lower horizontal line marks mean of CA (2.6 %).

## 4 Discussion

### 4.1 Temperature and rH are key factors to maintain cell viability in ALI experiments

Although a significant difference in viability between incubator and clean air control was detected, this difference is comparable to what is reported in other studies [26, 29, 30]. Recently, studies conducting experiments with ALI exposure systems highlighted the need to check if the used exposure system provides physiologically relevant conditions during exposure [21, 26, 31]. Often the existing exposure system needs to be modified prior to performing experiments to obtain realistic and reliable data [23]. For example, keeping temperature and rH at constant incubator-like conditions, e.g., 37 °C and 85 % rH, is of high importance for survival of cells during ALI exposure. A study by Zavala et al. showed that low temperature and low rH of the vehicle control (clean air) resulted in cytotoxic effects and low cell viability in ALI experiments [21]. Under these conditions, the potential effect of a test chemical could be masked by a high background toxicity due to poorly controlled exposure parameters. They also reported that only a small number of studies that conducted ALI experiments mention exposure parameters in detail. In our study, temperature and rH during experiments were at 38 °C and 65 % rH which approximates to 70 % rH at 37 °C, which is still below the target of 85 % rH reported in other studies [21, 31]. However, according to Zavala et al. a rH below 85 % could be practical since a lower rH at a certain temperature also means a lower dew point, which is the temperature at which water condenses in the air [21]. Further, a non-condensing atmosphere is desirable to protect downstream analysing devices [31]. The dew point for 38 °C and 65 % rH is approx. 30 °C. In our experiments, we observed small condensation in the outlet tubes of the exposure modules although the tubes were at temperatures of 36-37 °C, well above the calculated dew point. The condensation could have been excess humidity evaporating from the medium in the basal compartment of the exposure modules, as reported by Zavala et al. [21]. However, we observed no notably low volume of medium in the basal compartment of exposure modules after 1 h exposure. As mentioned, we installed a RH/T sensor upstream of each exposure module as a surrogate measurement for conditions at the insert-level inside the exposure modules. However, measurements upstream do not necessarily represent the conditions at the insert-level, where exposure of cells takes place. Guénette et al. observed a difference between measurements taken at the insert-level and downstream of the exposure module, suggesting not to rely on the surrogate measurements but to directly measure temperature and rH at the insert-level for better accuracy [31]. However, depending on the exposure system design, it could be difficult to place sensors at the insert-level for real-time monitoring of temperature and rH.

### 4.2 Delivery of test substances

Differences in concentration may arise from transport losses of test chemicals from chemical source to exposure module due to adsorption or reaction with pipes [17]. Since the pipes and tubes conducting the test chemicals in our experimental setup were made of stainless steel and inert tubing, we argue that adsorption or reaction was negligible. Guénette et al. established an ALI exposure system for testing of ozone with a feedback control loop that adjusts the ozone concentration at the source based on the downstream chemical analysis after the exposure module [31]. Kastner et al. used a downstream gas analyzer for formaldehyde and NO_2_ exposure [32]. When online monitoring of the test chemical cannot be achieved, different techniques for offline chemical analysis can be implemented. Bardet et al. analyzed the concentration of formaldehyde within the generation chamber using SPME on-fiber derivatization [33]. Persoz et al. put a passive sampler inside the exposure chamber and subsequently extracted the sampler after exposure for chemical analysis of formaldehyde [34]. A similar approach was chosen by Guénette et al. for delivery assessment of ozone [31]. A different approach is to perform deposition assays using reagents. Ritter et al. confirmed the delivery of NO_2_ onto the insert membrane using a reagent on the membrane that reacts with NO_2_ upon exposure which can be measured photometrically [35]. In our validation experiments, we performed a qualitative investigation of the delivery of NO_2_ onto the insert membranes by checking the inserts before and after exposure to 10 ppm NO_2_ for 1 h. After exposure, yellowish residuals distributed homogenously onto the insert membranes were observed which indicated deposition of NO_2_. Therefore, we reasoned that KEAA also was delivered to the exposure modules. Nevertheless, the effective concentration at cell level in the exposure module may have been different from the calculated concentration after dilution, which was based on the measurement of chemicals upstream of the exposure module.

### 4.3 Effects of gaseous KEAA fuel blend on A549 cells

In general, ALI experiments are considered more sensitive and more physiologically relevant for investigating inhalation toxicity of gaseous compounds compared to experiments using cells submerged in culture medium [15, 36, 37]. However, there are confounding factors during ALI exposure that may impair sensitivity of the exposure method, that might explain the few observed differences between control and treatments in this study.

First, the cells on the insert inside the exposure module may lose their air-liquid state when water droplets from condensing humidified air form a protective layer on the cell surface, effectively blocking direct interaction between test chemical and cells [16, 38]. In fact, we did observe condensation inside the outlet tubes of the exposure modules in our experiments, but no droplets on the trumpet-shaped inlets inside the exposure modules. Further, cells on inserts did not appear to be submerged or covered in droplets when examined under the microscope after exposure. Increasing temperature of the climatic chamber and water circuit system of the exposure modules helped reduce condensation [21, 26].

Second, a similar protective layer due to surfactants on the cell surface can reduce sensitivity to chemicals. During ALI cultivation, the air is a stimulus for cells to differentiate and produce surfactants. However, these physiological properties were only observed for A549 cells after a 6-day cultivation period at ALI [39]. Assumingly, the ALI cultivation period in our study (1 day) was too short to produce any significant protective surfactant layer on the cells. Cells cultivated at ALI are sensitive to external stress resulting from merely handling the cells. Thus, the ALI cultivation period during pre-incubation was kept to a minimum in this study.

Third, A549 cells lack the ability to form tight junctions [19, 39]. Without this barrier, biomarker, e.g., LDH, released from damaged cells on the insert should be able to diffuse from the apical side through the porous membrane into the basal medium. Studies investigating LDH content in apical washes of A549 cell layers and the basal medium showed that LDH content is higher in the basal medium [21, 40, 41]. Therefore, the lung model (A549) used in this study should be able to quantify biomarker of cytotoxicity.

Additional reasons why few effects in our study were observed could be that i) the exposure duration was short, ii) the recovery period too long or iii) the timepoint of endpoint investigation was too far apart from exposure.

The exposure duration of ALI experiments is limited to how long exposure parameters can be kept constant at incubator-like conditions during exposure. There are few studies that achieved this for up to 4 h [5, 26, 42]. However, keeping exposure parameters constant over a long time is difficult [17]. Therefore, many studies, including our, conducted ALI experiments up to 1 h of exposure [21, 32, 43, 44], which depicts a short-term acute exposure scenario. Despite the exposure duration limitation, we aimed to create a relevant exposure scenario. According to the GESTIS - International Limit Values for Chemical Agents [45] occupational exposure limits (OELs) of a given chemical must not be exceeded at the workplace. The OELs are used for workplace safety regulations and state average concentrations for both an 8 h work shift or a 15 min interval, in which the concentration can be higher than the 8 h value but must not occur more than 4 times during a work shift [46]. Since an 8 h exposure at ALI is technically difficult to accomplish, we designed the 1 h exposure duration to equal 4 times a 15 min interval with higher concentrations. The DFG OELs for 15 min exposure to the KEAA constituents are 400 (3-methylbutanone), 800 (ethanol), 400 (methyl acetate), 400 (ethyl acetate), 2000 (n-pentane) and 200 ppm (methanol) [45]. The concentrations of the dosed single KEAA constituents for the highest tested KEAA concentration (2000 ppm C_3.7_) based on measurements and calculations were 635.6, 396.6, 245.2, 196.8, 31.4 and 35.4 ppm, respectively. In this case, only the concentration of 3-methylbutanone (635.6 ppm) exceeded the OEL by a factor of approx. 1.6. Therefore, our data support the OELs of the KEAA constituents and moreover indicate no adverse effect to A549 cells when exposed to a mixture. This finding applies only to the exposure scenario tested in this study including the discussed drawbacks on the reliability of the effective concentration and is difficult to extrapolate to the real world since the investigated monoculture of A549 cells do not represent the complex *in vivo* situation.

## 5 Conclusion

Before experiments with the bio-hybrid fuel were performed, validation experiments were conducted to ensure high viability of A549 cells when exposed to humidified clean air. In our first experiments, viability of A549 cells exposed to humidified clean air was much lower than the incubator control and a high variance between replicates and experiments was observed. We identified temperature and rH as key contributors to the observed variance in the experiments. Therefore, we modified our exposure system to stabilize and monitor temperature and rH of the test gas more accurately in proximity to the cells. As a result, cell viability increased and variance in the data decreased.

In this study, A549 cells were exposed to a dynamic air flow of a freshly generated test gas containing a gaseous Ketone-Ester-Alcohol-Alkane (KEAA) bio-hybrid fuel blend. The concentration of KEAA was monitored online upstream of the Vitrocell exposure system. Data showed that constant and reproducible concentrations between experiments were achieved. However, the concentration upstream may differ from the effective concentration inside the exposure modules where exposure of cells takes place. A549 human lung cells were exposed at the ALI to 50-2000 ppm C_3.7_ gaseous KEAA for 1 h at 20 mL/min, 38 °C and 66 % rH. After 24 h recovery, there was a significant difference in LDH release and metabolic activity between clean air control and treatments in the mid-range of tested KEAA concentrations. However, no trend for concentration dependent change in LDH release or metabolic activity of cells was observed. This indicates that the significant differences observed might be artifacts due to a yet working but still improvable setup of the exposure device. For future experiments, focus on high relative humidity of the test gas and online downstream measurement of the effective concentration of chemicals is important to increase reliability of the data, making ALI experiments a suitable tool in early-stage bio-hybrid fuel research in the sense of green toxicology.

## 6 Acknowledgements

Dr. Richard Ottermanns for help and advice on statistical evaluation of data. Angelina Miller and Antonia Weltmeyer for helpful comments on the manuscript.

## 8 CRediT authorship contribution statement

**J. Daniel:** Conceptualization, Investigation, Validation, Formal analysis, Visualization, Writing – Original Draft Preparation. **A. Schönberger Alvarez:** Conceptualization, Investigation, Formal analysis, Visualization, Writing – Review & Editing. **P. te Heesen:** Investigation. **B. Lehrheuer:** Supervision, Funding Acquisition **S. Pischinger:** Supervision, Funding Acquisition. **H. Hollert:** Funding Acquisition, Writing – Review & Editing. **M. Roß-Nickoll:** Conceptualization, Supervision, Writing – Review & Editing. **M. Du:** Conceptualization, Validation, Supervision, Project Administration, Writing – Review & Editing.

## Notes

### Competing Interest Statement

The authors have declared no competing interest.

